# A novel Fc-optimized antibody-drug conjugate targeting CD7 for the therapy of T-cell acute lymphoblastic leukemia

**DOI:** 10.1101/2025.10.30.684608

**Authors:** Carina Lynn Gehlert, Denis Martin Schewe, Katja Klausz, Steffen Krohn, Dorothee Winterberg, Fotini Vogiatzi, Ammelie Svea Boje, Natalie Baum, Mona Könecke, Anja Lux, Falk Nimmerjahn, Regina Scherließ, Andreas Humpe, Martin Schrappe, Gunnar Cario, Thomas Valerius, Lars Fransecky, Anna Laqua, Monika Brüggemann, Roland Repp, Claudia Dorothea Baldus, Martin Gramatzki, Lennart Lenk, Christian Kellner, Matthias Peipp

## Abstract

While treatment for patients with T-cell acute lymphoblastic leukemia (T-ALL) has improved in the last decades, therapeutic options for patients refractory to standard therapy or with relapsing disease are limited. In particular, no immunotherapy option has been approved in T-ALL yet. Here, a novel dual antibody engineering approach for targeting CD7 was evaluated. The chimeric CD7 antibody chimTH69 was modified by Fc engineering to improve antibody-dependent cell-mediated cytotoxicity (ADCC) and antibody-dependent cellular phagocytosis (ADCP). In addition, it was conjugated to monomethyl auristatin E (MMAE), a microtubule-disrupting agent. The resulting Fc-optimized antibody-drug conjugate (ADC), designated chimTH69-DE-vcMMAE, showed a unique set of effector functions *in vitro*. It triggered ADCC by mononuclear cells at picomolar concentrations, mediated ADCP by macrophages and directly inhibited the growth of a panel of T-ALL cell lines by delivering the cytotoxic compound to induce G2 cell cycle arrest and apoptosis. In addition, due to its specific linker design, chimTH69-DE-vcMMAE demonstrated bystander killing activity against CD7-negative leukemia cells. In mice, CD7-directed therapy with chimTH69-DE-vcMMAE inhibited the growth of subcutaneous CCRF-CEM T-ALL xenografts. Moreover, chimTH69-DE-vcMMAE exerted strong antileukemic effects in a phase II-like patient-derived xenograft preclinical trial in pediatric and adult patients when applied in an experimental overt leukemia setting. ChimTH69-DE-vcMMAE induced minimal residual disease-negativity in one PDX model. These findings indicate that targeting CD7 with the novel Fc-optimized ADC is a potent strategy to trigger anti-leukemia responses and may open a novel therapeutic avenue for T-ALL treatment.

**Key Point:** A novel antibody drug conjugate targeting CD7 showed efficient anti-leukemia activity in preclinical models of T-ALL.

## Introduction

T-cell acute lymphoblastic leukemia (T-ALL) is an aggressive disease with complex molecular characteristics and accounts for approximately 15% of childhood and 25% of adult ALL cases [1-4]. Whereas in children approximately 80% of patients are cured, survival rates in adults only in selected subgroups are exceeding 50%. Therefore, novel therapeutic options are needed.

With the approval of blinatumomab and the antibody drug conjugate (ADC) inotuzumab-ozogamicin antibody-based immunotherapy has significantly improved the clinical outcome in B-ALL. Moreover, chimeric antigen receptor T (CAR-T) cell therapies have been established [5, 6]. Clinical trials investigating therapeutic antibodies in T-ALL are ongoing, including CD38 antibodies, but no antibody-based immunotherapy is yet approved for the treatment of T-ALL [7-9].

In T cell neoplasia, CD7 represents a promising target antigen. CD7 is strongly expressed in all T-ALL subtypes including pro-T, pre-T, cortical T and mature T-ALL. Importantly, also patients with early T-cell precursor (ETP) ALL display strong CD7 expression [10, 11]. Moreover, CD7-expression has been described on leukemia initiating cells of T-ALL [12]. In the hematopoietic system, CD7 is expressed by subsets of T and myeloid progenitor cells, natural killer (NK) cells, and the majority of mature T cells in the peripheral blood [13]. Hematopoietic stem cells and 5 to 20% of peripheral T cells lack CD7 expression, which may become relevant in therapeutic settings targeting CD7 to maintain immune function required for the prevention of infections [13-15].

Based on these characteristics, CD7 has been evaluated as a target antigen for different antibody-based approaches in previous studies. In earlier studies our group demonstrated that the murine IgG1 antibody TH69 was effective in eliminating T-ALL xenografts in mice by Fc-dependent modes of action [16]. Due to its high internalization capacity, CD7 was analyzed as a target structure for immunotoxins and ADC [15, 17-20]. Although technically challenging, CD7-directed CAR T cells have been established and exhibited first promising results in early clinical trials in T-ALL patients further validating CD7 as a feasible target structure for therapeutic intervention [21-25].

In this study, a novel Fc-optimized ADC, chimTH69-DE-vcMMAE, was designed and its anti-leukemia activity was evaluated in preclinical models of T-ALL. ChimTH69-DE-vcMMAE was effective via a dual mode of action - depleting T-ALL cells by activating immune effector cells, and by inducing cell death via the conjugated cytotoxic payload MMAE. Our *in vivo* experiments proved efficient anti-leukemia activity and showed significant long-term survival after CD7-ADC therapy. These data further validate CD7 as a promising target for ADC in T-ALL.

## Methods

### Cell lines

CEM, Jurkat, MOLT-16, HSB-2, P12, Karpas-45 and Nalm-6 cells were obtained from the DSMZ (German Collection of Microorganisms and Cell Cultures); CHO-S cells were purchased from Thermo Fisher Scientific. All cell lines were cultured as recommended. CHO-FcγRIIa cells and BHK-FcγRIIIa cells were cultivated as described [26-29].

### Human effector cell separation / primary leukemia cells / PDX samples

Experiments were approved by the Ethics Committee of the medical faculty (Kiel, Germany), in accordance with the Declaration of Helsinki (D552/17). All participants gave written informed consent. Peripheral blood mononuclear cells (PBMCs) from healthy donors were isolated as previous described [26]. Macrophages were generated as already specified [30].

### Production of antibodies and ADC

The chimeric and Fc optimized CD7 antibody (chimTH69-DE) was generated using the variable light (VL) and heavy chain (VH) sequences of the murine antibody TH69, established in our laboratory [16]. VL and VH sequences were cloned into modified human antibody light chain (LC) or IgG1 heavy chain (HC) expression vectors, which were derived from pSecTag2/HygroC. The HC sequence contained nucleotide exchanges for the S239D/I332E amino acid exchanges [31]. The antibody was produced and purified as previously described [30]. The ADC chimTH69-DE-vcMMAE was generated by conjugation of MMAE via disulfide bonds and the enzyme-cleavable linker mc-vc-PABC to chimTH69-DE by a commercial provider (Cfm Oskar Tropitzsch GmbH).

### Cell viability

Growth inhibitory effects were analyzed with 1×10^4^ cells per well in 96-well plates treated for 96 h with serial dilutions of the indicated antibodies and the MTT proliferation assay (Cell Proliferation Kit I, Roche) following the manufacturer’s recommendations.

### Flow cytometry

To investigate antigen binding, quantification of expression levels, induction of apoptosis, cell cycle analysis and bystander activity immunofluorescence analyses were performed as described in *Supplementary Methods*.

### Cytotoxicity and phagocytosis assays

Antibody-dependent cellular phagocytosis (ADCP) and antibody-dependent cell-mediated cytotoxicity (ADCC) were analyzed as previously described [30, 32].

### *In vivo* experiments

Animal experiments were approved by the local authorities. In all *in vivo* approaches, female NOD.Cg-Prkdc^SCID^Il2rg^tm1Wjl^/SzJ (NSG) mice (Charles River Laboratories) were used. 1×10^6^ CEM cells were subcutaneously (s.c.) injected into the right flank or 0.5×10^6^ T-ALL patient-derived xenografts (PDX) cells were injected intravenously into the tail vein of mice. Animals were treated intraperitoneally (i.p.) with a dose of 1 mg/kg body weight of antibody (chimTH69-DE-vcMMAE or chimTH69-DE).

### Statistical analyses

Data were statistically analyzed using suitable tests with GraphPad Prism 10 Software (GraphPad). Significance was accepted with *P*<0.05.

Further methodological details are available in the *Supplementary Methods*.

## Results

### CD7 expression levels in T-ALL and generation of a Fc-optimized CD7-specific antibody drug conjugate

First, CD7 surface expression was quantified on the cell surface of eight T-ALL PDX samples (Table 1), five T-ALL patients and a panel of T-ALL cell lines. T-ALL PDX samples showed comparable expression levels compared to primary T-ALL cells and cell lines. CD7 antigen surface expression was in the range of 2×10^4^ to 2×10^5^ molecules per cell described as specific antibody binding capacity (SABC) that was in most cases higher than CD7 expression on non-malignant T cells (Figure 1, Table 1).

**Table 1.**
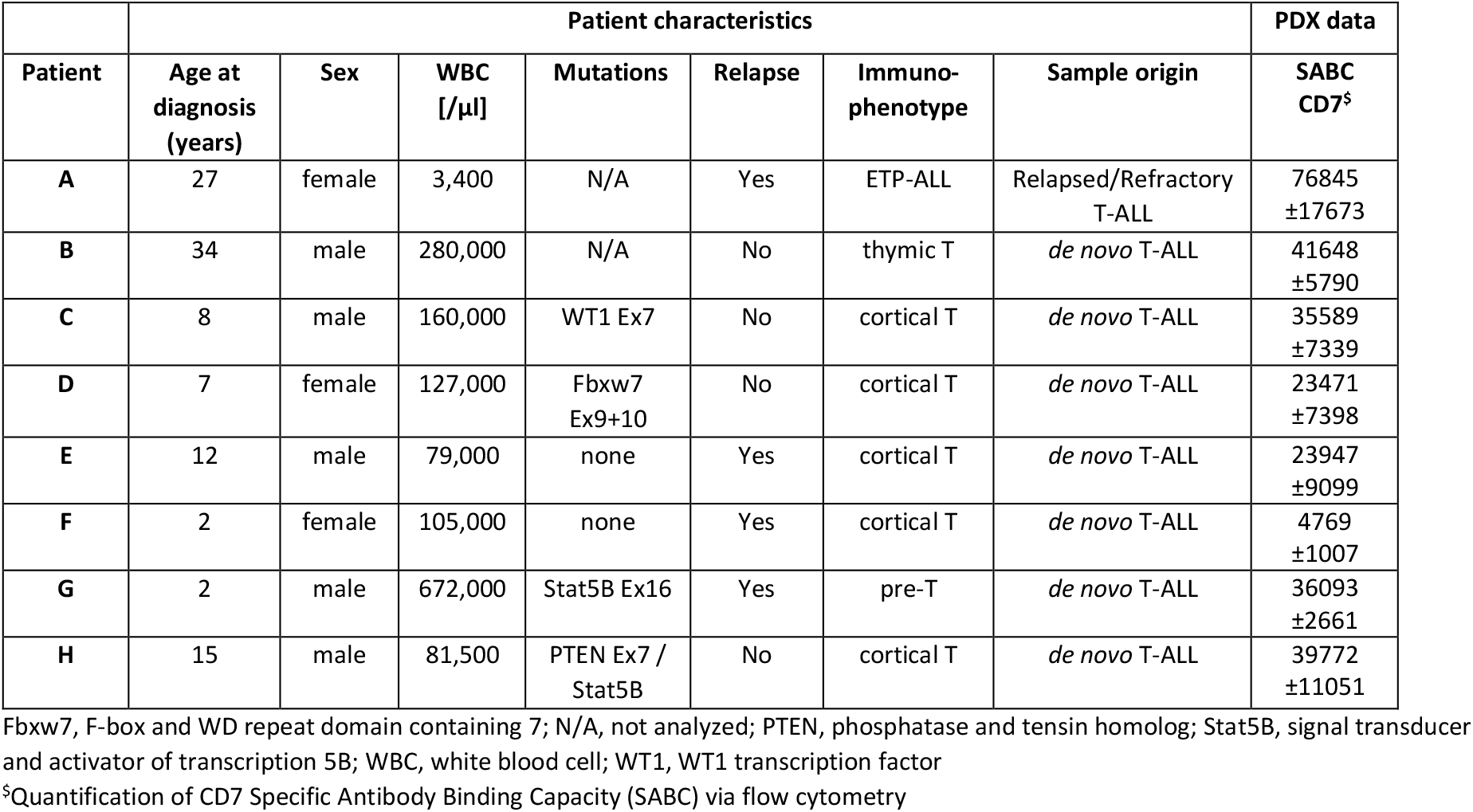
Characteristics of T-ALL patients with samples used for the randomized preclinical phase 2-like xenograft study.

**Figure 1.**
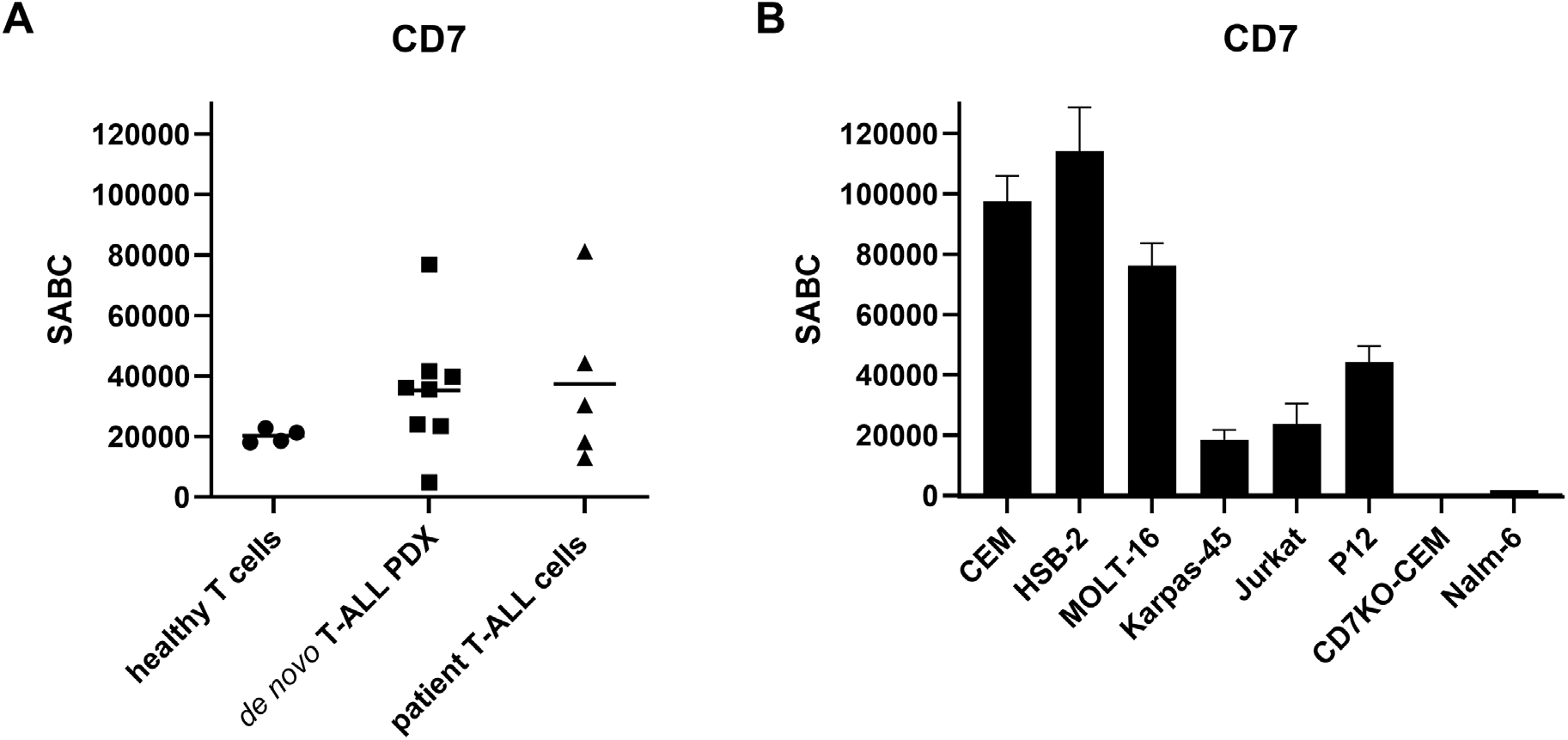
Quantification of CD7 cell surface expression levels. The expression of CD7 was quantified as the specific antigen binding capacity (SABC) on **A.** T cells from healthy donors, T-ALL PDX samples, primary patient cells and **B**. T-ALL cell lines in comparison to a CD7-knockout CEM cell line (CD7KO-CEM) and the CD7^-^ BCP-ALL cell line Nalm-6. Mean values ± SEM of n=4 (T-ALL PDX samples)/ n=3 (T-ALL cell lines) independent experiments.

To target CD7, an antibody engineering approach was used. Our chimeric CD7 antibody carrying the variable regions of the murine TH69 antibody [16] was optimized for enhanced Fcγ receptor (FcγR) binding and its ability to trigger ADCC and ADCP by introducing two amino acid substitutions (S239D/I332E; DE-variant; [33]) in the Fc part. The resulting Fc-optimized CD7 antibody chimTH69-DE was conjugated to the microtubule-disrupting agent MMAE via an enzymatically cleavable linker (mc-vc-PABC). This resulted in the ADC chimTH69-DE-vcMMAE with a drug-to-antibody ratio (DAR) of 3.2 MMAE-molecules per antibody (supplemental Figure 1A/B). The Fc-optimized ADC and the unconjugated antibody showed similar binding capacity to CD7^+^ T-ALL cell lines, whereas no binding to CD7^-^ cells was observed (supplemental Figure 1C). The concentration dependent binding of chimTH69-DE-vcMMAE and chimTH69-DE revealed EC_50_ values in the low nanomolar range (chimTH69-DE: EC_50_ = 1.38 nM; chimTH69-DE-vcMMAE: EC_50_ = 3.09 nM). The ADC had a slightly reduced binding avidity, which suggested a minor negative impact of the MMAE conjugation on CD7 binding (supplemental Figure 1D). Moreover, both chimTH69-DE-vcMMAE and chimTH69-DE bound cells stably transfected with either FcγRIIa or FcγRIIIa (supplemental Figure 1C).

### ChimTH69-DE-vcMMAE induces G2-cell-cycle arrest and apoptosis in CD7^+^ T-ALL cell lines

We next characterized the direct growth inhibitory activity of chimTH69-DE-vcMMAE against different CD7^+^ T-ALL cell lines. Representative cell lines, which were classified to different T-ALL subtypes by their immunophenotype (Pro-, pre-, mature- and cortical-T-ALL; [34]) and which demonstrated varying CD7 surface expression levels, were evaluated (supplemental Table 1; Figure 1B). ChimTH69-DE-vcMMAE showed significant dose-dependent growth inhibition in all CD7^+^ cell lines with IC_50_ values at low nanomolar concentrations (0.21 - 1.18 nM; supplemental Table 1; Figure 2A), while CD7^-^ cell lines were not affected. The unconjugated antibody was not effective in this setting (Figure 2A). An almost linear correlation between the CD7 expression on the cell surface (SABC) and maximal growth inhibition after ADC treatment was observed (Figure 2B). Cell lines with high CD7 expression (SABC >76,000), such as HSB-2 and MOLT-16, showed the highest maximal growth inhibition of up to 98.7%, whereas in Karpas-45 cells (SABC = 18,465) cell viability was reduced by 45.2%. Yet, CEM cells with the highest CD7 expression level (SABC = 97,525) showed a maximal growth inhibition rate of 82.9%. These data suggest that the antigen expression level is apparently not the only critical factor determining the susceptibility to chimTH69-DE-vcMMAE but certainly plays an important role. To further demonstrate the specificity of chimTH69-DE-vcMMAE, blocking experiments with the parental murine antibody mTH69 were performed. As expected, no reduction in cell viability was detected after CD7 blocking, which proves a strict target antigen-specific cytotoxic effect of chimTH69-DE-vcMMAE (Figure 2C).

**Figure 2.**
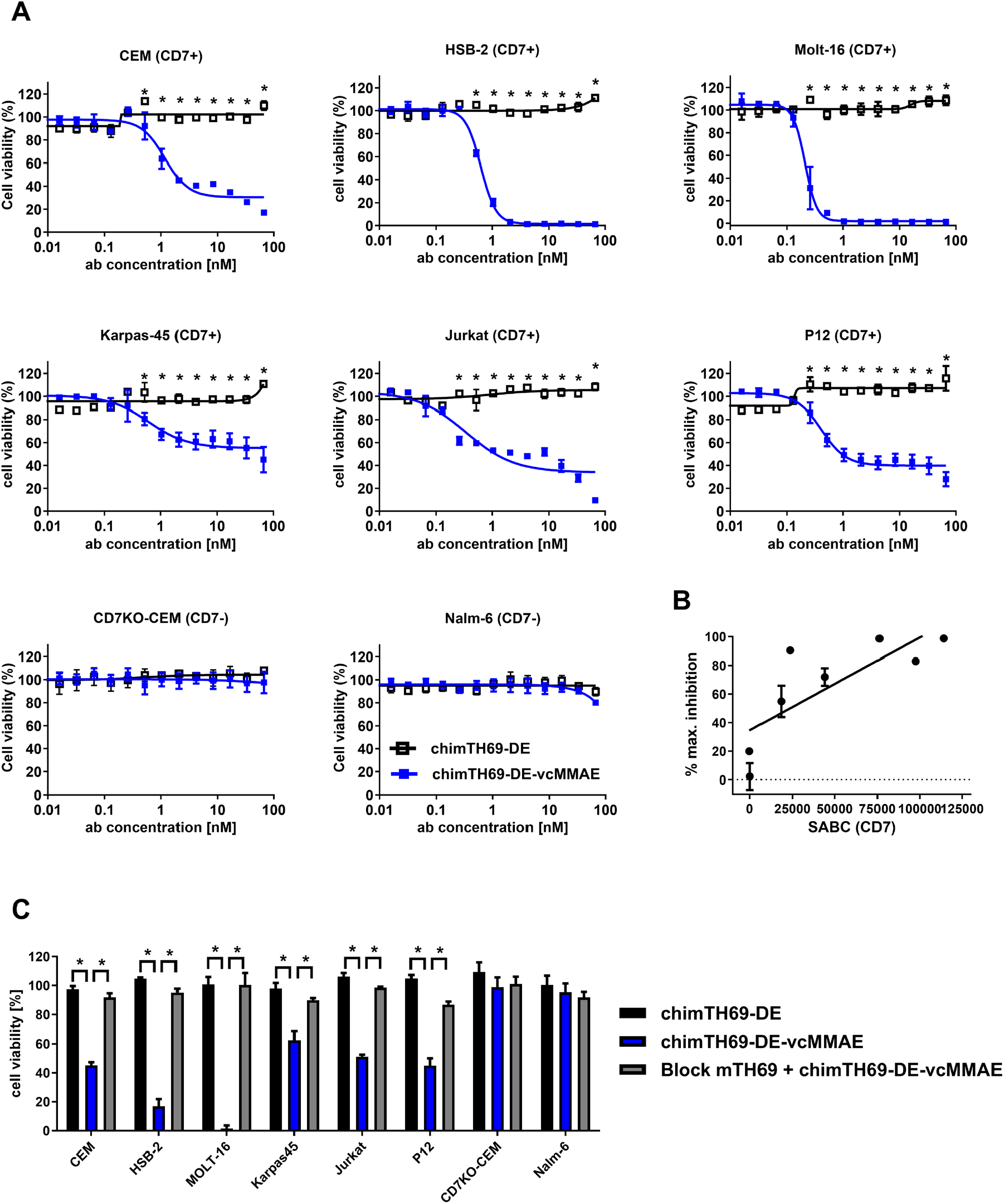
Direct anti-proliferative effects of chimTH69-DE-vcMMAE against target antigen expressing cell lines. **A**. CD7^+^ T-ALL cell lines, the CD7^-^ B cell line Nalm-6 and CD7-knockout CEM cells (CD7KO-CEM) were treated with increasing antibody (ab) concentrations of chimTH69-DE-vcMMAE (blue filled symbols) or chimTH69-DE (black open symbols) for 96 h and the cell viability was tested by the MTT assay. Mean values ± SEM of n=3 independent experiments, * *P*<0.05 chimTH69-DE-vcMMAE vs. chimTH69-DE, two-way ANOVA with Bonferroni post-test. **B**. The maximal inhibition of the cell viability in percent shows a linear correlation to the CD7 Specific Antibody Binding Capacity (SABC) of the depicted cell lines (Pearson correlation r = 0.79, p = 0.02). Mean values ± SEM of n=3 independent experiments. **C**. As a control for antigen-specific effects, cells were pretreated with a 10-fold molar excess of the parental murine TH69 antibody (Block mTH69) for 30 minutes to mask the CD7 epitope (blockade) and were incubated afterwards with chimTH69-DE-vcMMAE (2 nM) for 96 h. The cell viability was tested by the MTT assay. Mean values ± SEM of n=3 independent experiments, * *P*<0.05, one-way ANOVA with Bonferroni post-test.

MMAE acts by potently inhibiting tubulin polymerization. Therefore, the induction of cell cycle arrest and apoptosis by chimTH69-DE-vcMMAE was analyzed. CEM cells were treated with chimTH69-DE-vcMMAE and the cell cycle was analyzed by flow cytometry. ChimTH69-DE-vcMMAE treatment led to a significant increase of cells in G2/M phase and to a significant reduction of cells in G1 (Figure 3A, B). In addition, the sub-G1 fraction was higher in the chimTH69-DE-MMAE treated cells, which indicates the induction of cell death (Figure 3A). None of these effects were observed in untreated controls, chimTH69-DE treated cells, or when the cells were preincubated with the parental murine antibody mTH69 (Figure 3A,B).

**Figure 3.**
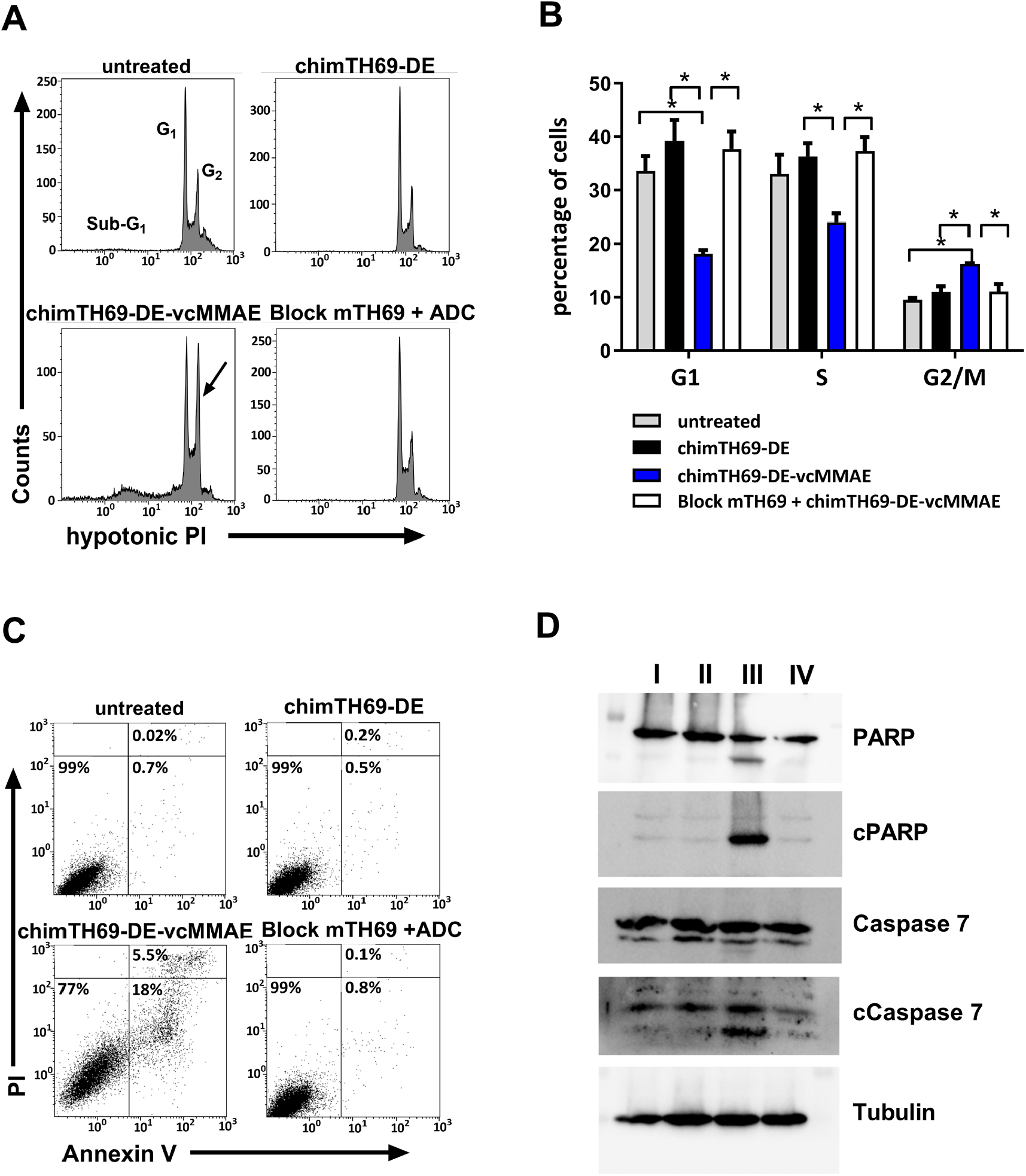
ChimTH69-DE-vcMMAE mediates apoptosis and G2-cell cycle arrest in CD7^+^ cell lines. **A-C.** CEM cells were treated with chimTH69-DE or chimTH69-DE-vcMMAE (3 nM) or were left untreated (medium) for 72 h. For target antigen blocking CEM cells were preincubated for 30 minutes with the parental murine TH69 (30 nM) antibody (Block mTH69). **A**. Representative histograms of flow cytometry analysis of hypotonic PI staining of the cell nucleus and cell cycle analysis. **B**. Quantitative analysis of cell cycle phases. Mean values ± SEM of n=4 independent experiments, * *P*<0.05. One-way ANOVA with Bonferroni post-test. **C**. Representative histograms of flow cytometry analysis of Annexin V and PI staining of early and late apoptotic cells. **D**. Representative images of western blot analysis of pro-apoptotic proteins PARP and Caspase 7 and their cleavage products (c) in CEM cells after treatment with chimTH69-DE (II), chimTH69-DE-vcMMAE (III) (3 nM) or medium (untreated, I) for 72 h. For target antigen blocking, cells were preincubated for 30 minutes with the murine parental antibody (mTH69, 30 nM, IV). Tubulin served as a loading control.

To further evaluate the type of cell death, cell surface exposure of phosphatidylserine, PARP cleavage and caspase 7 cleavage was analyzed. ChimTH69-DE-vcMMAE treatment showed early (Annexin V^+^ / PI^-^) and late (Annexin V^+^ / PI^+^) apoptotic cells (Figure 3C). Preincubation with the parental antibody mTH69 prevented apoptosis induction in chimTH69-DE-vcMMAE treated cells (Figure 3C), confirming a strict target antigen-dependent mode of action. The data were further confirmed by demonstrating cleavage of PARP and caspase 7 in chimTH69-DE-vcMMAE treated cells (Figure 3D). Together, these data prove that chimTH69-DE-vcMMAE is able to act by cell cycle arrest and apoptosis induction.

### Bystander anti-tumor activity of chimTH69-DE-vcMMAE against CD7^-^ cell lines

After the intracellular cleavage of the valine-citrulline linker by cathepsins, the active metabolite (free MMAE) has been demonstrated to be permeable, to cross the cell membrane and to be able to diffuse into neighboring cells, like antigen-negative tumor cells (antigen escape) or stroma cells in the tumor microenvironment. The killing of antigen-negative neighbor cells is called bystander effect or bystander killing and has been suggested as a clinically relevant mode of action by recently approved ADC, such as trastuzumab-deruxtecan [35, 36].

To examine the bystander killing capacity of chimTH69-DE-vcMMAE, CD7^-^ (CEM cells with CD7-knockout) and CD7^+^ cells (CEM cells) were cultivated in co- or in single culture in the presence or absence of chimTH69-DE-vcMMAE. The number of Annexin-V^+^ cells in each cell population was quantified via flow cytometry. As tumor antigen escape models, we used a CD7-knockout T-ALL cell line (CD7KO-CEM, Figure 4A) and CD7^-^ B-ALL Nalm-6 cells (Figure 4B, supplemental Table 1). A significant number of apoptotic CD7^+^ CEM cells in co-culture and in mono-culture in the presence of the chimTH69-DE-vcMMAE was observed (Figure 4A + 4B). In CD7KO-CEM cells (Figure 4A) and Nalm-6 cells (Figure 4B), the induction of apoptosis was detected only in co-culture with CD7^+^ cells (CEM), but not in mono-culture settings. These data demonstrate that chimTH69-DE-vcMMAE is able to trigger bystander killing, potentially extending its mode of action.

**Figure 4.**
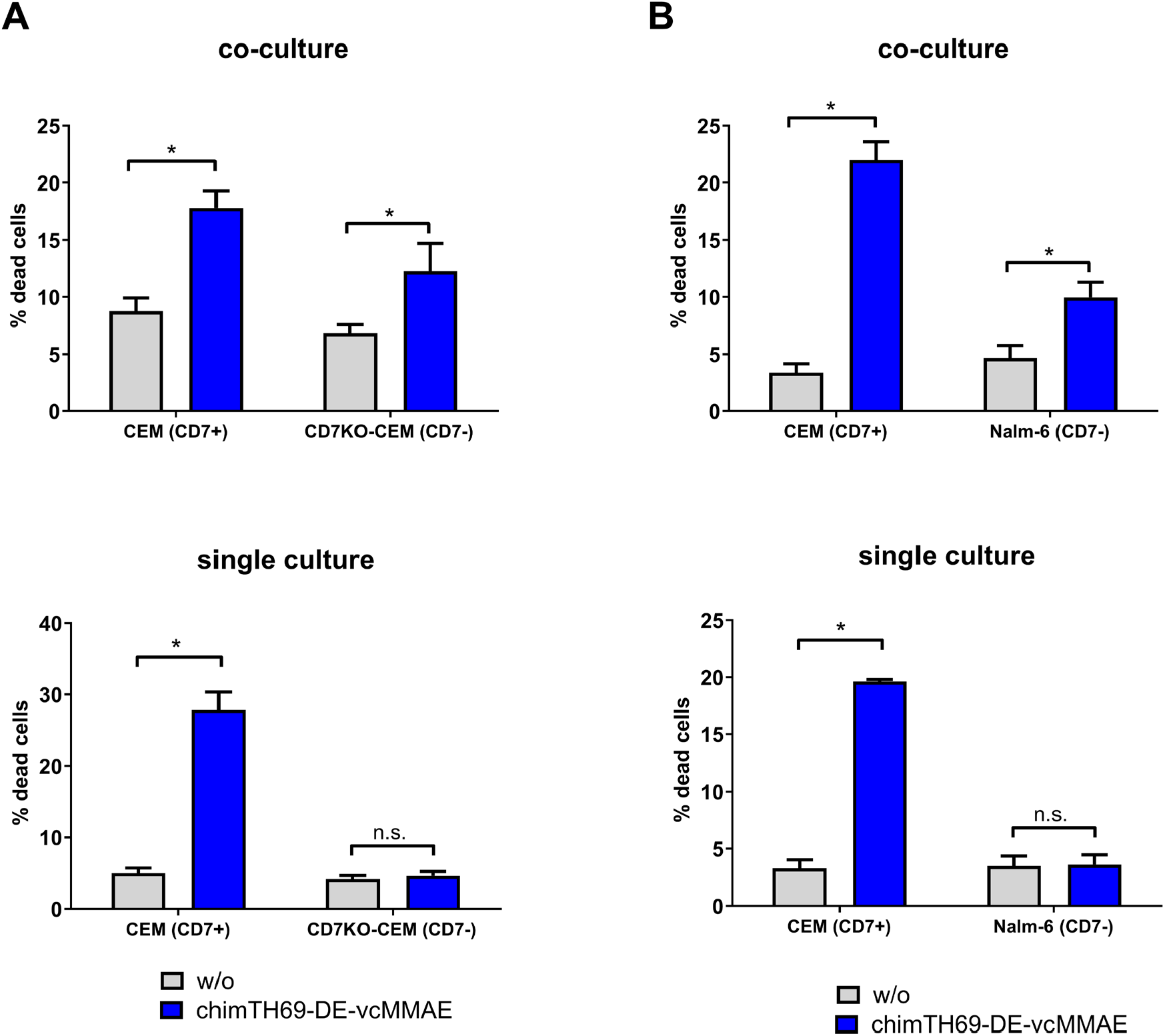
ChimTH69-DE-vcMMAE triggers bystander killing of CD7^-^ ALL cells in co-culture with CD7^+^ target cells. **A+B.** CD7^+^ (CEM cells) and CD7^-^ cells (**A**. antigen escape surrogate cells: CD7KO-CEM cell line or **B**. BCP-ALL cell line Nalm-6) were treated with 10 nM chimTH69-DE-vcMMAE or left untreated (medium, w/o) under co-culture conditions (ratio 1:1) or in single culture. The amount of Annexin V^+^ cells was analyzed via flow cytometry after 72 h. Mean values ± SEM of n=3 (Nalm-6)/n=5 (CD7KO-CEM) independent experiments, * *P*<0.05, n.s. not significant, one-way ANOVA with Bonferroni post-test.

### The Fc-optimized CD7-ADC effectively triggers ADCC and ADCP

ChimTH69-DE-vcMMAE is based on an IgG1 Fc backbone carrying two amino acid exchanges for improved Fc receptor binding also present in the clinically approved antibody tafasitamab [37, 38]. To investigate, if the optimized CD7-specific ADC was able to mediate Fc-dependent effector functions, the capacity to trigger ADCC and ADCP was evaluated. Efficient ADCC of different CD7^+^ T-ALL cell lines was mediated by chimTH69-DE-vcMMAE with PBMC as effector cells (Figure 5A). At saturating concentrations, no significant differences between chimTH69-DE-vcMMAE and chimTH69-DE were observed. However, at low concentrations, chimTH69-DE-vcMMAE showed slightly reduced target cell lysis compared to chimTH69-DE. This could be explained by the observed differences in CD7 binding between chimTH69-DE-vcMMAE and chimTH69-DE (supplemental Figure 1D). Nevertheless, chimTH69-DE-vcMMAE showed improved ADCC activity compared to the non-engineered chimTH69-wt antibody (Figure 5A). To confirm that the detected tumor cell lysis is mediated through activation of effector cells and not through direct cytotoxic effects of MMAE during the short assay duration, control experiments in the absence and presence of effector cells were performed (Figure 5B). As expected, no tumor cell lysis was detected without effector cells, whereas efficient lysis of different T-ALL cell lines was shown in the presence of PBMC (Figure 5B). The capacity of chimTH69-DE-vcMMAE to trigger ADCP was analyzed with different T-ALL cell lines (CEM, HSB-2, MOLT-16) and monocyte-derived macrophages as effector cells. The Fc-optimized ADC, as well as the unconjugated antibodies (chimTH69-DE and chimTH69-wt), mediated ADCP of various T-ALL cells (Figure 5C).

**Figure 5.**
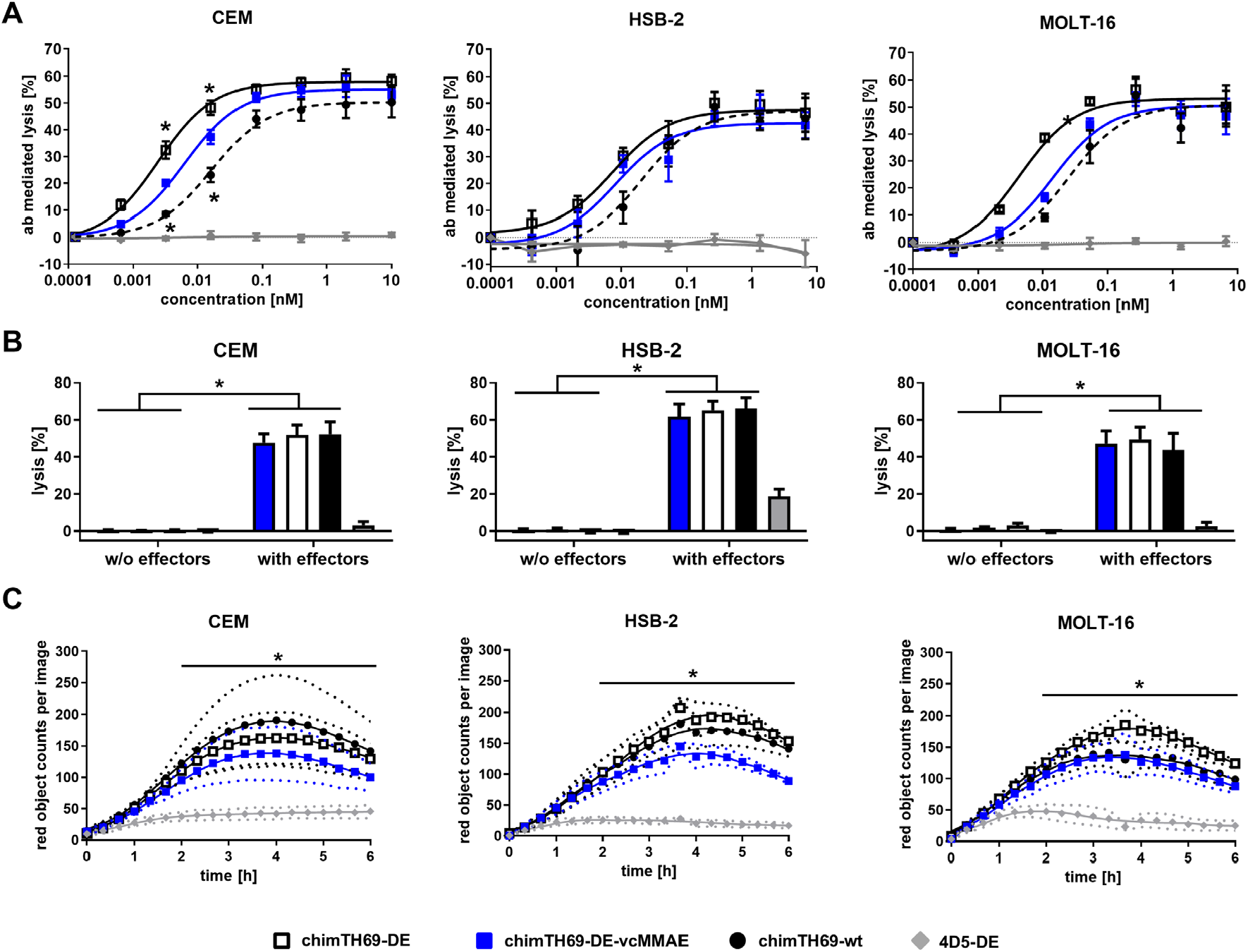
ChimTH69-DE-vcMMAE triggers ADCC and ADCP. **A.** ADCC of CD7^+^ tumor cell lines (CEM, HSB-2 and MOLT-16) was analyzed in standard chromium release assays with increasing antibody concentrations and PBMCs of healthy donors at an effector-to-target cell (E:T) ratio of 40:1. Antibody-mediated tumor cell lysis was tested for chimTH69-wt, chimTH69-DE or chimTH69-DE-vcMMAE and was compared to the 4D5-DE control antibody. Mean values ± SEM of n=3 independent experiments, * *P*<0.05 chimTH69-DE-vcMMAE vs. chimTH69-DE or chimTH69-wt, two-way ANOVA with Bonferroni post-test. **B**. ADCC of CD7^+^ tumor cell lines (CEM, HSB-2 and MOLT-16) was performed in standard chromium release assays at an antibody concentration of 1 µg/ml (6.67 nM) in the presence or in the absence of effector cells (PBMC). Mean values ± SEM of n=3 independent experiments, * *P*<0.05 w/o effector cells vs. with effector cells for chimTH69-DE, -wt, -DE-vcMMAE, two-way ANOVA with Bonferroni post-test. **C**. Phagocytosis was measured for 6 h by high-throughput fluorescence microscopy. CD7^+^ tumor cells (CEM, HSB-2 and MOLT-16) labelled with a pH-sensitive red-fluorescent dye were incubated at an E:T ratio of 1:1 with macrophages and 10 µg/ml of the indicated antibodies. Phagocytosis is depicted as the red object counts per image. Mean values ± SEM of n=3 independent experiments, * *P*<0.05 chimTH69-DE-MMAE, chimTH69-DE or chimTH69-wt vs. 4D5-DE, two-way ANOVA with Holm-Sidak’s post-test (CEM) or Mixed-effect model (REML) with Holm-Sidak’s post-test (HSB-2 and MOLT-16).

In summary, chimTH69-DE-MMAE is able to mediate direct cytotoxic effects via the conjugated MMAE and to trigger Fc-mediated effector functions.

### ChimTH69-DE-vcMMAE shows therapeutic activity in T-ALL cell line xenograft and PDX models

In the next sets of experiments, we investigated the therapeutic activity of chimTH69-DE-vcMMAE in xenograft mouse models of T-ALL. First, chimTH69-DE-vcMMAE was tested in the well-established CEM T-ALL cell line xenograft model. CEM cells were injected s.c. in immunodeficient NSG mice and animals were either treated with 1 mg/kg of chimTH69-DE-vcMMAE or chimTH69-DE or left untreated (Figure 6A). Animals that were treated with chimTH69-DE-vcMMAE showed a significant reduction in tumor growth (Figure 6A). Antineoplastic activity was detectable with the unconjugated antibody chimTH69-DE probably due to a functional myeloid effector cell compartment; however, its therapeutic efficiency was significantly lower than that of chimTH69-DE-vcMMAE (Figure 6A). No differences in body weight were detected between the treatment groups indicating that application of chimTH69-DE-vcMMAE was generally well tolerated in this setting (supplemental Figure 2).

**Figure 6.**
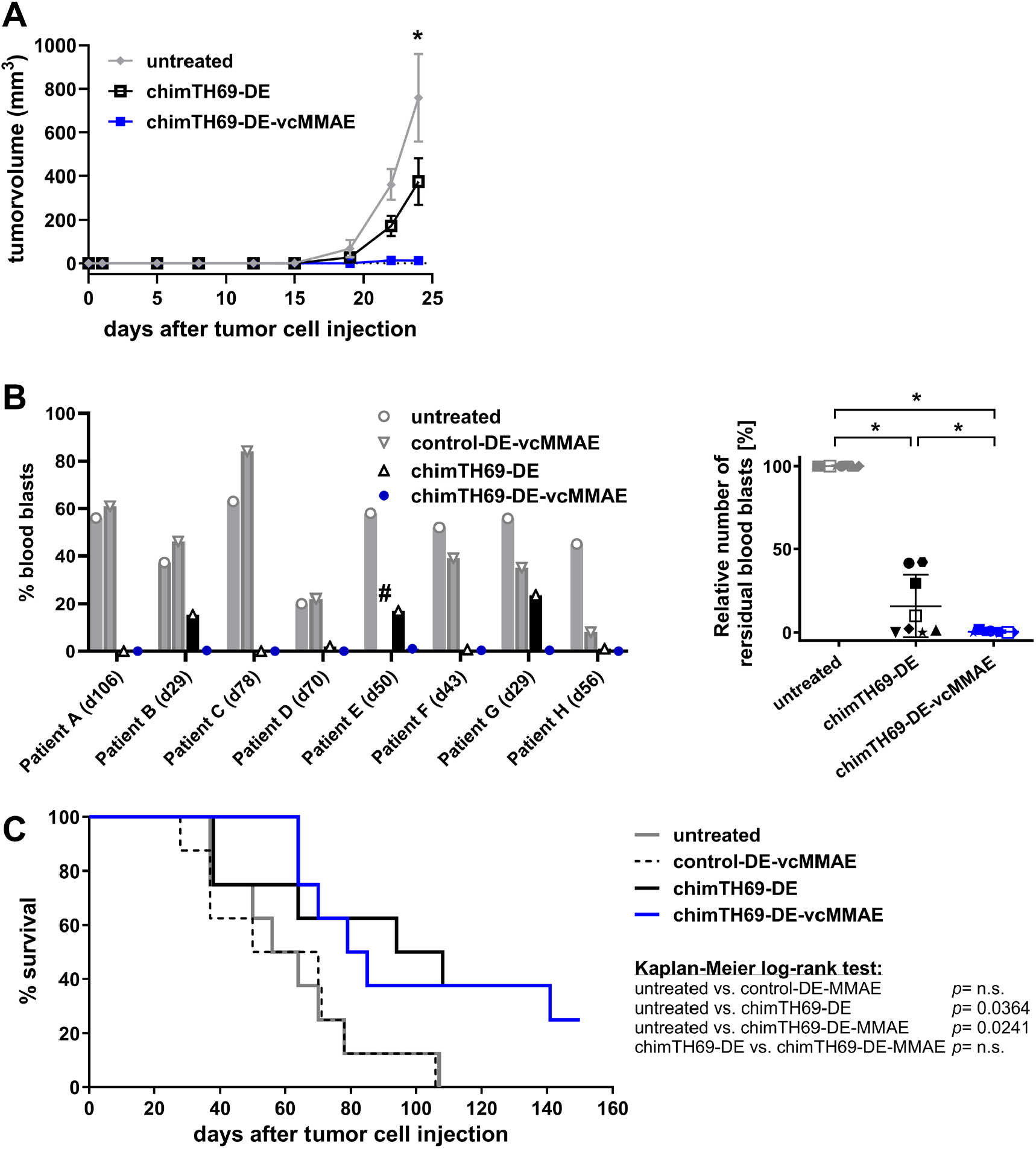
*In vivo* anti-tumor efficacy of chimTH69-DE-vcMMAE in T-ALL xenografts in mice. **A.** 24 h after subcutaneous (s.c.) injection of CEM cells in NSG mice, animals were treated twice weekly with chimTH69-DE-vcMMAE (5 mice per group), chimTH69-DE (10 mice per group) or with vehicle control (phosphate buffered saline (PBS); untreated; 10 mice per group). Tumor volume was calculated by regular caliper measurement of s.c. tumors. Tumor volumes are depicted until day 24 when the first control mouse was taken out of the experiment. * *P*<0.05, untreated vs. chimTH69-DE-vcMMAE; two-way ANOVA with Bonferroni post-test. **B + C**. Phase 2-like preclinical study in an overt leukemia setting with eight T-ALL PDX samples injected into NSG mice. Per patient, four PDX were established and the animals received therapy with chimTH69-DE-vcMMAE, chimTH69-DE, control-DE-vcMMAE or were left untreated. **B**. Analysis of blood blasts by flow cytometry at time points when the control animals had a mean blast load of 48 %. # = Control-DE-vcMMAE treated mouse of Patient E died on day 28 without a sign of leukemia. * *P*<0.05, n.s. not significant; one-way ANOVA with Holm-Sidak’s post-test. **C**. Survival was analyzed using the Kaplan-Meier method and log-rank statistics, n.s. not significant.

We next analyzed the efficacy of chimTH69-DE-vcMMAE in a preclinical phase 2-like PDX study using eight T-ALL-PDX samples (one sample generated from a relapsed/refractory (r/r) and seven PDX models from previously untreated patients (de novo T-ALL) [39, 40]). PDX cells from pediatric and adult patients (patient A - H; Figure 1B, Table 1) were intravenously injected into four NSG mice per patient, and antibody therapy was started when 1% human blasts were detected in peripheral blood (overt leukemia study) as published previously [39, 40]. Animals were treated with 1 mg/kg of chimTH69-DE-vcMMAE, chimTH69-DE, a similarly designed control ADC targeting an irrelevant antigen (control-DE-vcMMAE), or were left untreated. PDX-ALL engraftment was traced by detection of hCD45^+^/hCD7^+^ cells in peripheral blood (PB). Comparative PB analysis was conducted when one of the four corresponding PDX mice showed clinical signs of leukemia or >50% ALL-PDX cells in the PB [41]. Whereas control-DE-vcMMAE had no anti-leukemic efficacy, both chimTH69-DE-vcMMAE and chimTH69-DE significantly reduced the leukemic burden in PDX animals. All PDX models significantly responded to CD7-directed therapy. For all PDX mice treated with chimTH69-DE-vcMMAE no blast counts were observed, whereas in three mice of the chimTH69-DE treatment group significant blast numbers were detectable. The therapeutic effect of chimTH69-DE-vcMMAE was significantly more pronounced compared to chimTH69-DE (Figure 6B). Accordingly, chimTH69-DE-vcMMAE therapy resulted in a significantly prolonged median survival in comparison to mice treated with the control ADC or untreated animals (Figure 6C). Likewise, animals receiving therapy with the unconjugated CD7-antibody showed a significant prolongation of median survival compared to control groups (Figure 6C). Interestingly, no significant difference in the survival was detected between the two CD7-specific therapies (chimTH69-DE-vcMMAE and chimTH69-DE) (Figure 6C). The experiment was terminated on day 150. At this time point, all control animals were sacrificed because of high leukemic blast counts in the peripheral blood, while 2 of 8 mice of both CD7-specific treatment groups survived until the end of the experimental period without developing signs of leukemia (patient A and patient C, respectively). To analyze the depth of remission in surviving animals, the MRD status was determined in the bone marrow of these mice. Surviving mice of patient A treated with chimTH69-DE-vcMMAE or chimTH69-DE were low positive. Interestingly, the surviving animal xenografted with cells from patient C and receiving chimTH69-DE-vcMMAE treatment was MRD negative, whereas the surviving animal of the same patient treated with chimTH69-DE was MRD positive. (supplemental Table 2).

In summary, the Fc-optimized ADC is able to trigger Fc-mediated effector functions, mediates direct cytotoxicity through the conjugated MMAE and is able to act on neighboring cells via bystander killing. Importantly, chimTH69-DE-vcMMAE showed efficient anti-tumor effects in two mouse models, including one model of adult and r/r T-ALL. Therefore, chimTH69-DE-vcMMAE represents a promising new immunotherapeutic agent for treating T-ALL, warranting further preclinical and clinical investigation.

## Discussion

CD7 is an attractive target antigen for immunotherapy, since most T cell neoplasias display high expression [8]. In line with previous studies, we demonstrated stable CD7 expression in T-ALL patients [8, 10, 11]. The absence of CD7 on hematopoietic stem cells and subpopulations of T cells and NK cells [13, 14] may maintain certain immune functions and ensure the reconstitution of the T cell compartment after CD7-directed therapy [15]. This aspect was confirmed in first clinical studies with CD7-directed CAR T cells, with T cell and NK cell counts recovering to normal levels in about half of the patients [21-23].

A key property of CD7 is its rapid internalizing capacity, making it an ideal target for immunotoxins and ADC [15-19]. Accordingly, chimTH69-DE-vcMMAE was effective against T-ALL cell lines with different CD7 expression levels by triggering G2/M cell cycle arrest and induction of apoptosis at low nanomolar concentrations, in line with characteristics of approved MMAE-containing ADCs such as brentuximab vedotin and polatuzumab vedotin [36, 42]. Although the uptake of chimTH69-DE-vcMMAE is strictly antigen-dependent, it is capable to act on CD7^-^ neighboring cells by its bystander activity. Bystander killing plays an important role in the therapeutic activity of ADC, such as trastuzumab deruxtecan against tumors with heterogeneous target expression [35, 43]. Whether bystander activity is beneficial for CD7-targeting in patients is speculative, but may be warranted to allow targeting the tumor microenvironment / stem cell niche of leukemic stem cells and may prevent CD7-negative relapse as observed in early CAR T cell trials [23].

It is well established that many clinically approved ADC mediate Fc effector functions [35, 44, 45]. In this study, chimTH69-DE-vcMMAE by its engineered Fc domain was capable to induce significant FcγR-mediated effector mechanisms. Whether Fc receptor binding / Fc-mediated effector mechanisms are beneficial for the antitumor activity of ADC or contributes to toxic side effects is still discussed controversially [35, 36, 46]. The clinically approved Fc-optimized ADC belantamab mafodotin triggers strong FcγR-mediated effector mechanisms [47]. Barely any changes in the proliferation and survival of FcγR-expressing cells were observed after belantamab mafodotin treatment *in vitro* and interestingly an increased number of FcR-bearing macrophages in treated mice were observed [47]. Likewise, our Fc-optimized chimTH69-DE-vcMMAE did not show significant target independent FcR-mediated cytotoxicity against PBMCs and macrophages in first *in vitro* analyses (unpublished data). Whether the benefit of improved Fc-mediated effector functions outweighs potential side effects needs further evaluation.

In a subcutaneous xenograft mouse model chimTH69-DE-vcMMAE demonstrated significant reduction in tumor growth. Also the unconjugated antibody chimTH69-DE as previously described for its murine counterpart, from which it was derived [16], showed some therapeutic activity. However, the activity of chimTH69-DE was significantly reduced in comparison to chimTH69-DE-vcMMAE. Since NSG mice lack functional NK cells, probably murine myeloid effector cells were responsible for the antitumor effects of the unconjugated CD7-antibody. This is consistent with previous studies demonstrating that human IgG1 antibodies bind to mouse FcγRIV expressed on mouse myeloid cells [48-50]. Together, these data indicate that the Fc-optimized chimTH69-DE antibody induces Fc-mediated effector mechanisms in subcutaneous T-ALL xenograft models, whereas the Fc-engineered CD7-ADC additionally acted via the conjugated MMAE payload. The therapeutic activity of chimTH69-DE-MMAE was further confirmed in PDX models of pediatric and adult T-ALL patients. A significant reduction of blast cell count was achieved in all PDX models with chimTH69-DE-vcMMAE treatment and even MRD-negativity was reached in one PDX model. Interestingly, a significant response was also achieved in the PDX model derived from a patient with ETP-ALL, indicating that anti-leukemia activity may also be expected in these high-risk T-ALL patients. Interestingly, leukemia cells in relapsing animals still expressed CD7 and were sensitive to chimTH69-DE-vcMMAE treatment, indicating that optimized treatment regimens may further improve the therapeutic activity.

In contrast to the s.c. xenograft model also the unconjugated antibody chimTH69-DE showed significant therapeutic efficacy. Reasons for differences in the two *in vivo* studies can be an interplay of various parameters. In general, differences in the pharmacokinetic properties of ADC and unmodified antibodies in the murine system may underestimate the activity of ADC since fast metabolism of ADC has been described especially in NSG mice [51]. Further studies in Fc receptor knockout mice or depletion of murine effector cells will be required to further dissect the relative contribution of the different modes of action of our ADC *in vivo* [52-54].

In summary, chimTH69-DE-vcMMAE showed significant activity *in vitro* and *in vivo*, which indicates that CD7 targeting with the novel Fc-optimized ADC is a potent strategy to trigger anti-leukemia responses. In clinical settings the combination of ADC such as gemtuzumab ozogamicin or belantamab mafodotin with chemotherapy or immunomodulatory drugs showed synergistic potential and may also represent further strategies to achieve long-lasting therapeutic activity in a higher proportion of patients treated with CD7-ADC [55-57]. Since CD7-directed therapy with chimTH69-DE-vcMMAE will likely result in significant reduction of malignant and non-malignant T cells this type of therapy may allow patients to proceed safely to allogeneic stem cell transplantation. The safety and efficacy of the proposed concept of an Fc-engineered ADC targeting CD7 needs to be studied in clinical trials and may open up a novel therapeutic avenue for patients with T cell neoplasia.

## Acknowledgements

We thank Britta von Below, Anja Muskulus, Irene Pauls and Gabriele Riesen for excellent technical assistance.

The manuscript contains data in partial accomplishment of the requirements for a thesis by C.L.G at the Faculty of Mathematics and Natural Sciences of the Kiel University.

This work was funded by the *Deutsche Jose Carreras Leukämie-Stiftung* (DJCLS 20 R/2021, to C.K. and M.P.). Contributing groups are in part supported by the Deutsche Forschungsgemeinschaft (DFG, German Research Foundation)—project number 444949889 (KFO 5010 Clinical Research Unit “CATCH ALL” to D.S., M.B., L.L., K.K., G.C., M.S. C.D.B. and M.P.).

## Authorship Contributions

C.L.G. designed and performed the experiments and analyzed data; L.L., K.K., M.K., D.W., S.K., N.B., F.V., A.S.B. performed experiments and analyzed data; A.L., F.N., R.S., A.H., M.S., G.C., T.V., L.F., M.B., C.D.B., M.G., L.L., D.M.S. provided essential reagents, cell lines, PDX material and patient information; M.P., C.K. and C.L.G. wrote the manuscript; C.K. and M.P. initiated and designed experiments and supervised the study. All authors discussed and approved the manuscript.

