## Supplementary material for "A novel Fc-optimized antibody-drug conjugate targeting CD7 for the therapy of T-cell acute lymphoblastic leukemia": Suppl Methods + Data

### Supplemental methods

**Generation of CD7-negative T-ALL cell line:** CD7-knockout in CCRF-CEM cells was generated via CRISPR/Cas9 technology. Briefly, 2.5 x $10^6$ CEM cells were transfected with 1.5 µM Alt-R® S.p. Cas9 Nuclease V3 (Integrated DNA Technologies) and 4.4 µM guideRNA (gRNA; published sequence was *de novo* synthesized by Synthego[1]) via the MaxCyte (STX) Scalable Transfection System, as described by the manufacturer. CD7-knockout CEM cells were isolated by cell-sorting the CD7-negative cell fraction with a FACSAria (BD) automated cell sorter.

**Western blot analysis:** Apoptosis induction was analyzed with cells treated for 72 h with the indicated antibodies or left untreated. 50 µg protein of cell lysate were loaded on 12 % Tris–acrylamide gels under reducing conditions and were blotted to PVDF membranes according to standard procedures. For apoptosis detection PARP (#9542), cleaved-PARP (#5625), Caspase 7 (#12827), cleaved Caspase 7 (#8438) antibodies (Cell Signaling Technology) were added to a final dilution of 1:5000. Detection of tubulin (#ab18257, Abcam) served as a loading control. As a secondary antibody the HRP-conjugated anti rabbit IgG (#7074, Cell Signaling Technology) was used at a final dilution of 1:5000. Blots were finally analyzed using a chemoluminescent substrate (Pierce, Thermo Fisher Scientific) and a ChemiDoc imaging system (Biorad).

**Flow cytometry:** Immunofluorescence analyses were performed on a Navios flow cytometer (Beckman Coulter) and analyzed with Kaluza Analysis software (Beckman Coulter).
For Investigation of monoclonal antibody and ADC binding, cells were incubated with 50 µg/ml of the indicated antibodies (1 h on ice). For concentration dependent binding analysis, cells were treated with antibodies at varying concentrations (1 h on ice). Binding was detected using secondary anti-human-kappa-FITC antibody (SouthernBiotech; 30min on ice).
Quantification of CD7 expression levels were examined with the CD7-specific antibody (mTH69) and the QIFI Kit (Agilent Dako) according to the manufacturers' protocols.
For the analysis of apoptosis induction, cells were treated for 72 h with the indicated antibodies or left untreated. Early and late apoptotic cells were stained using AnnexinV-APC/PI (Biolegend). Cell cycle analysis was measured using a hypotonic PI solution as described previously [2] and was analyzed using FlowJo Software (BD, Becton Dickinson).
To analyze the bystander anti-tumor activity, CD7-positive cells were labelled with the membrane dye DiI (Thermo Fisher Scientific) and cultured for 72 h at 37 °C in co-culture with unlabeled CD7-negative cells (ratio 1:1) in the presence of the ADC or left untreated. Apoptotic/necrotic cells (dead cells) were analyzed after 72 h with AnnexinV-APC (Biolegend) staining and cell populations were separated via gating DiI-positive and DiI-negative cells.

**Xenograft models:** In subcutaneous xenografts CEM T-ALL cells were injected at day 0 into animals and treatment was administered on days +1, +5, +8, +12, +15, +19 and +22 intraperitoneally (i.p.). The tumor volume was calculated by regular caliper measurement of subcutaneous tumors. All animals were sacrificed when the tumor volume was larger than 1500 $mm^3$.
In PDX xenografts T-ALL PDX cells were injected into animals. Experiments were performed as randomized preclinical phase II-like study using eight different T-ALL PDX. Randomized phase II-like xenograft trials can represent the heterogeneity of patient populations [3-5]. The therapy was started when 1% blasts were detected in the peripheral blood (overt leukemia model [4]) and antibodies were injected on day +1, +3, +6, +10, +13 as described previously [4] and every 7 days thereafter. Leukemic engraftment was studied via flow cytometric detection of human CD45+/CD7+/CD38+ cells versus murine CD45+ cells in the peripheral blood. Mice were sacrificed upon detection of >75% leukemic blasts or upon signs of leukemic engraftment. Isolation of bone marrow and minimal residual disease (MRD) analysis were performed as previously described [4-6]. Briefly, BM samples of mice that survived the experiments were analyzed for the persistence of MRD by quantitative PCR for patient-specific immunoglobulin/T-cell receptor rearrangements (IgTR) or by digital droplet PCR quantifying human albumin when no IgTR markers were available[6].

### Supplemental figures

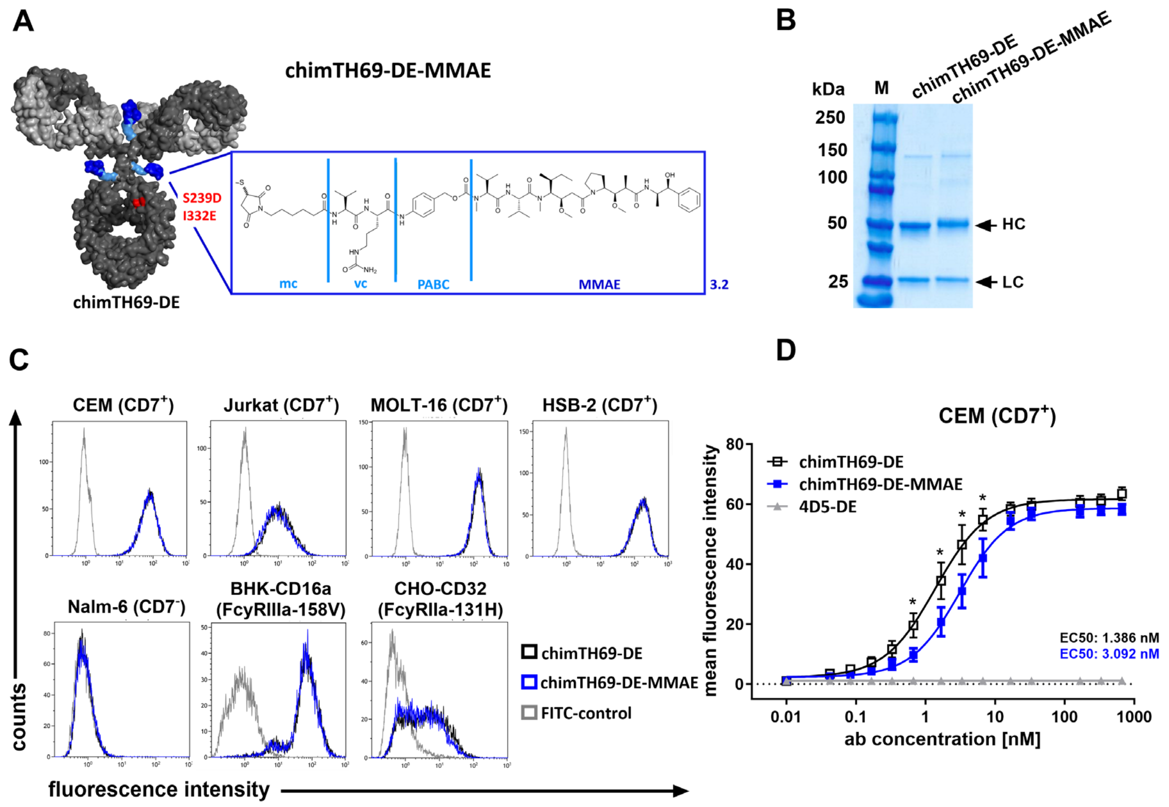


**Supplemental figure 1 Generation of the CD7-specific antibody drug conjugate A**. Schematic illustration of chimTH69-DE-MMAE, an Fc-optimized CD7-specific antibody drug conjugate. The chimeric IgG1 antibody directed against CD7 with two amino acid exchanges (S239D and I332E; red) in the Fc-part was linked to the cytotoxic compound monomethyl auristatin E (MMAE; dark blue) via an enzymatically cleavable linker (mc-vc-PABC; light blue). The drug to antibody ratio was 3.2 MMAE-molecules per antibody. **B.** Validation of purity and mass of the ADC and the unconjugated antibody by SDS-PAGE under reducing conditions followed by Coomassie blue staining. **C.** Antigen specific binding of chimTH69-DE-vcMMAE and chimTH69-DE was tested via flow cytometry on different CD7$^+$ T-ALL cell lines (CEM, Jurkat, HSB-2 and MOLT-16) and on the CD7$^-$ BCP-ALL cell line Nalm-6. The binding of the optimized Fc-part to different FcγR was also tested via flow cytometry on two different stably transfected cell lines (BHK-CD16a and CHO-CD32). Data show representative histograms of n=3 experiments. **D.** Binding analysis of chimTH69-DE-MMAE and chimTH69-DE in a concentration dependent manner compared to a control antibody (4D5-DE) was tested on CD7$^+$ cell line CEM via flow cytometry. Mean values ± SEM on n=3 independent experiments, * $P$<0.05 chimTH69-DE-vcMMAE vs. chimTH69-DE, Two-way ANOVA with Bonferroni post-test.

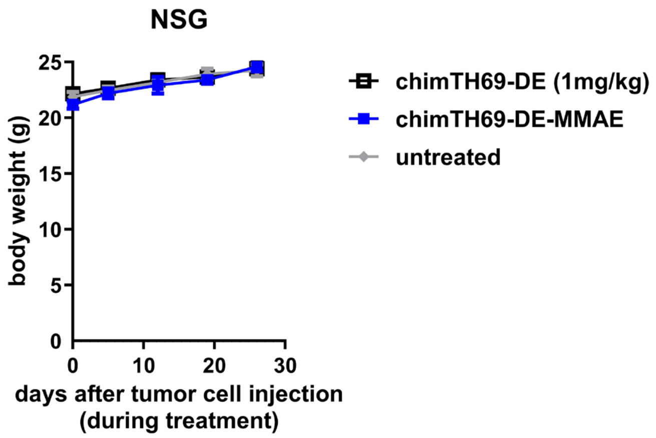


**Supplemental figure 2 Body weight of NSG mice during treatment.** 24 h after subcutaneous injection of CEM cells in NSG mice, animals were treated twice weekly with chimTH69-DE-vcMMAE, chimTH69-DE or with vehicle control (phosphate buffered saline (PBS); untreated). Body weight was observed during treatment period until day 26.

### Supplemental tables

**Table 1 CD7⁺ T-ALL cell lines and CD7-negative BCP-ALL cell line Nalm-6 and CD7-knockout T-ALL cell line CEM.** Quantification of CD7 Specific Antibody Binding Capacity (SABC) on different T-ALL cell lines assigned to the T-ALL subtypes: Pro-T-ALL, Pre-T-ALL, Cortical T-ALL and Mature T-ALL [7]. $IC_{50}$ values and maximal inhibition of the cell viability in percent measured in MTT-assay. Mean values ± SEM on n=3 independent experiments.

| | | SABC (CD7) | $IC_{50}$ [nM] | max. Inhibition [%] |
|---|---|---|---|---|
| **CCRF-CEM** | Pre- T-ALL | 97,525 ± 4,924 | 1.18 ± 1.16 | 82.9 ± 1.3 |
| **HSB-2** | Pre- T-ALL | 114,151 ± 8,422 | 0.63 ± 1.03 | 98.7 ± 1.2 |
| **MOLT-16** | Mature- T-ALL | 76,227 ± 4,246 | 0.21 ± 1.06 | 98.7 ± 1.4 |
| **Jurkat** | Mature- T-ALL | 23,623 ± 3,432 | 0.34 ± 1.34 | 90.4 ± 0.6 |
| **P12/ICHIKAWA** | Cortical- T-ALL | 44,129 ± 3,143 | 0.41 ± 1.15 | 71.9 ± 6.2 |
| **Karpas 45** | Pro- T-ALL | 18,465 ± 1,887 | 0.61 ± 1.43 | 54.9 ± 11.1 |
| **Nalm-6** | BCP-ALL | 204 ± 12 | - | 19.9 ± 1.1 |
| **CD7KO-CEM** | CD7 knockout T-ALL | 85 ± 23 | - | 2.3 ± 9.3 |

**Table 2 Minimal residual disease (MRD) measurement in isolated DNA of bone marrow samples of surviving animals.** n.d. = not determined; n.m. = not measured

| Patient | untreated | control-DE-vcMMAE | chimTH69-DE | chimTH69-DE-vcMMAE |
|---|---|---|---|---|
| A | n.m. due to leukemic engraftment | n.m. due to leukemic engraftment | low positive | low positive |
| B | n.d. | n.d. | n.d. | n.d. |
| C | n.m. due to leukemic engraftment | n.m. due to leukemic engraftment | positive | negative |
| D | n.d. | n.d. | n.d. | n.d. |
| E | n.d. | n.d. | n.d. | n.d. |
| F | n.d. | n.d. | n.d. | n.d. |
| G | n.d. | n.d. | n.d. | n.d. |
| H | n.d. | n.d. | n.d. | n.d. |
